# Integrating Genome-Wide Association Analysis, Functional Annotation and Regulatory Genomics for the Prioritization of Candidate Variants Associated with Grain Yield in Upland Rice

**DOI:** 10.64898/2026.07.10.737825

**Authors:** Agnes Cardoso da Cruz, Rosana Pereira Vianello, Paula Arielle Mendes Ribeiro Valdisser, Luíce Gomes Bueno, Claudio Brondani

## Abstract

Grain yield is a highly complex quantitative trait in rice, resulting from the interaction of multiple genetic, physiological and environmental factors. Although genome-wide association studies (GWAS) have successfully identified loci associated with grain yield, translating statistical associations into biologically meaningful candidate variants remains a major challenge, particularly for variants located in regulatory regions. This study aimed to identify genomic variants associated with grain yield in upland rice and to develop an integrative framework for functionally prioritizing candidate variants through the combination of genome-wide association analysis, functional annotation and regulatory genomics. A panel of 252 accessions from the Brazilian Rice Core Collection was phenotyped for grain yield and genotyped with 35,763 single nucleotide polymorphism (SNP) markers. GWAS identified 29 SNPs significantly associated with grain yield, including 16 variants located within or near annotated genes and 13 located in intergenic regions. The identified candidate genes were involved in signal perception, metabolite transport, amino acid and energy metabolism, hormone biosynthesis, protein turnover, RNA processing and disease resistance, highlighting the polygenic architecture of grain yield. Functional characterization of the intergenic regions revealed enrichment of *cis*-regulatory elements recognized by transcription factors associated with hormonal signaling, drought response, carbon metabolism, photosynthesis and reproductive development, indicating that regulatory variation represents an important component of grain yield determination. By integrating GWAS signals, candidate gene annotation, *cis*-regulatory element characterization and the physical proximity between SNPs and *cis*-regulatory elements, an integrative prioritization strategy identified seven intergenic SNPs as the most promising candidates for functional validation. Together, these findings establish a robust framework for discovering, prioritizing and functionally validating regulatory variants, bridging the gap between statistical associations and biological function while providing a rational strategy for translating GWAS discoveries into molecular breeding of complex traits.

## INTRODUCTION

Grain yield is the primary objective of rice breeding programs, particularly in rainfed cropping systems, where water deficit, high temperatures, and low soil fertility often limit the expression of cultivars’ yield potential (Fernando et al., 2025). As a complex quantitative trait controlled by multiple genes and strongly influenced by the environment, its expression results from the interaction between various physiological processes and regulatory networks that coordinate growth, development, and adaptation to environmental conditions.

Genome-wide association studies (GWAS) have become a fundamental tool for identifying loci associated with complex quantitative traits, enabling the exploration of genetic variability in germplasm collections and natural populations. In rice, numerous studies have identified genomic regions associated with grain yield; however, the biological interpretation of GWAS signals of these associations remains a major challenge in association genomics (Ashfaq et al., 2023). In most cases, the identified SNPs serve as statistical markers, while the identification of potentially causal variants and underlying molecular mechanisms remains limited.

This challenge is particularly evident for variants located in non-coding regions of the genome. Recent evidence demonstrates that a large proportion of phenotypic variation in quantitative traits can result from alterations in *cis*-regulatory elements that modulate the intensity, timing, and spatial specificity of gene expression, without directly modifying the protein-coding sequence (Joly-Lopez et al., 2020; Zhu et al., 2024). Despite this growing recognition, the contribution of SNPs located in regulatory regions remains underexplored in rice GWAS studies, particularly in germplasm adapted to rainfed cultivation environments in tropical regions.

The integration of GWAS, functional annotation of candidate genes, and characterization of regulatory regions represents a promising strategy to enhance the interpretative power of association studies (Qi et al., 2024). Beyond identifying loci associated with the phenotype, this strategy enables a biological evidence framework to prioritize candidate variants for functional validation, bridging the gap between association genomics and functional genomics—as well as their application in molecular breeding programs.

In this context, this study aimed to identify genomic variants associated with grain yield in a panel of 252 accessions from the Embrapa rice core collection—genotyped with 35,763 SNPs—and to develop an integrative framework for the functional prioritization of these variants by combining GWAS, functional annotation of candidate genes, and regulatory genomics. To this end, candidate genes were functionally characterized, intergenic regions enriched in *cis*-regulatory elements were analyzed, and the proximity between SNPs and *cis*-regulatory elements was incorporated as an additional criterion to prioritize variants with greater biological plausibility and potential application in functional validation studies and molecular breeding.

## MATERIAL AND METHODS

Genotyping of the 252 accessions was performed using the genotyping-by-sequencing (GBS) technique, following the methodology described by Pantalião et al. (2016). After aligning the reads to the rice reference genome Os-Nipponbare-Reference-IRGSP-1.0/MSU7, SNPs were identified using a minimum allele frequency of 0.01, an inbreeding coefficient of 0.9, and a minimum locus coverage of 0.1. Population structure was estimated using the software Structure (Pritchard et al., 2000), with the most probable number of subpopulations determined by the method of Evanno et al. (2005) using the Structure Harvester program (Earl & von Holdt, 2012).

Genome-wide association studies (GWAS) was performed using Tassel v.5.2.44 software (Bradbury et al., 2007) and a mixed linear model (MLM), with the population structure matrix (Q) as a fixed effect and the kinship matrix (K)—estimated via the Identity-by-State (IBS) algorithm—as a random effect. Significant associations between SNPs and yield were identified using an FDR threshold of ≤ 0.05 (Storey, 2002). Functional annotation of significant SNPs was performed using the RiceVarMap (https://ricevarmap.ncpgr.cn/v3/) and Rice Genome Annotation Project (RGAP, https://rice.uga.edu/) platforms. For SNPs located in intergenic regions, sequences between adjacent genes were extracted using the Os-Nipponbare-Reference-IRGSP-1.0/MSU7 reference genome. *cis*-Regulatory elements within the intergenic regions containing the identified SNPs were characterized using the PlantCARE platform (Lescot et al., 2002). Priority was given to *cis*-elements associated with hormonal responses, water deficit, carbon metabolism, reproductive development, and grain yield regulation.

## RESULTS AND DISCUSSION

A total of 35,763 SNPs were identified across the set of accessions studied, with an average density of 95.83 SNPs/Mbp. Population structure analysis revealed two main genetic groups (K = 2), reflecting both the evolutionary history and the domestication and breeding processes of the evaluated accessions. The first group consisted predominantly of cultivars and improved lines, whereas the second was composed mainly of traditional varieties. This separation aligns with studies on rice germplasm collections, in which materials subjected to successive selection cycles exhibit reduced genetic variability due to the fixation of alleles favorable to agronomic traits, whereas traditional varieties maintain greater genetic diversity and serve as important reservoirs of novel alleles for breeding programs (Huang et al., 2022; Singh et al., 2024).

A genome-wide association analysis identified 29 SNPs significantly associated with grain yield, reinforcing the highly polygenic nature of this trait in rice. Unlike qualitative traits controlled by a few genes of large effect, yield results from the interaction of multiple physiological processes related to vegetative growth, reproductive development, energy metabolism, assimilate transport, hormonal regulation, and adaptation to environmental conditions. The distribution of SNPs across different chromosomes and functional classes, along with the identification of 16 candidate genes with distinct biological functions, confirmed that yield is determined by the integrated action of multiple molecular pathways—a result consistent with previous genome-wide association studies in rice (Table 1). The identified candidate genes span diverse functional categories related to the crop’s agronomic performance. Notable examples include genes involved in cellular signal perception and transduction, such as members of the receptor-like kinase (RLK) family; metabolite transport, represented by *OsALMT7*; and amino acid metabolism, such as threonine synthase (LOC_Os01g49890), whose role in essential amino acid biosynthesis makes its association with vegetative growth, nitrogen use efficiency, and grain filling biologically plausible (Zou et al., 2015; Heng et al., 2018).

**Table 1.** Coding-region SNPs associated with grain yield in upland rice and their corresponding candidate genes. SNP nomenclature: the number following the letter “S” indicates the chromosome, followed by the SNP genomic position.

| SNP | Alleles * | Gene (ID / annotation) |
| --- | --- | --- |
| S1_27476179 | C/T | LOC_Os01g48000 – Receptor-like kinase (RLK) |
| S6_8962815 | G/A | LOC_Os06g15779 – ALMT malate transporter |
| S8_5457694 | G/T | LOC_Os08g09420 – OsFBX266 (F-box) |
| S3_17070977 | A/C | LOC_Os03g29950 – G6PDH |
| S2_23154369 | T/G | LOC_Os02g38290 – Cytochrome P450 |
| S6_24557073 | C/A | LOC_Os06g41090 – OsFTIP1 (FT-Interacting Protein 1) |
| S4_34336548 | C/G | LOC_Os04g57670 – PPR (pentatricopeptide) |
| S7_17400233 | T/G | LOC_Os07g29600 – C3HC4 Ring zinc-finger |
| S4_28494263 | C/A | LOC_Os04g47912– 5' UTR variant |
| S12_1869646 | G/A | LOC_Os12g04410 - 5' UTR variant |
| S9_20349326 | G/C | LOC_Os09g34920 – Glycosyl hydrolase 29 |
| S1_28654895 | A/C | LOC_Os01g49890 - Threonine synthase |
| S3_27301283 | C/G | LOC_Os03g48030 - Integral membrane HPP family protein |
| S7_17404811 | T/G | LOC_Os07g29610 - PQ loop repeat domain |
| S8_8951845 | G/A | LOC_Os08g14850 - Disease resistance protein |
| S11_17596780 | A/C | LOC_Os11g30290 - PHF5-like protein domain |

\* Alleles are presented as Reference allele/Alternative allele.

The results also highlighted genes related to energy metabolism and the control of cellular redox balance, such as glucose-6-phosphate dehydrogenase (G6PDH), as well as genes from the cytochrome P450 superfamily involved in phytohormone biosynthesis and degradation (Peng et al., 2024; Nagasawa et al., 2013; Xu et al., 2014). Furthermore, proteins from the F-box, RING zinc-finger, PPR, and PHF5-like families participate in ubiquitination and RNA processing—processes essential for development, metabolism, and environmental adaptation (Jain et al., 2007; Wang et al., 2022; Meng et al., 2024; Liu et al., 2025). Genes related to disease resistance (LOC_Os08g14850) and flowering (*OsFTIP1*) were also identified, reinforcing that grain yield depends on the coordination between growth, development, and environmental response (Song et al., 2017; Hu et al., 2021). Although distinct from classic grain yield genes such as *Gn1a, DEP1, IPA1, GS3, GW2*, and *Ghd7*, these genes converge on the same physiological networks responsible for determining grain yield, representing new candidates for functional validation and molecular breeding.

One of the most significant findings of this study was the identification of 13 grain yield-associated SNPs located in intergenic regions (Table 2). Recent evidence demonstrates that a large proportion of phenotypic variation in quantitative traits stems from alterations in regulatory regions capable of quantitatively modulating gene expression (Li et al., 2022; Manzo et al., 2025). In this context, the identified SNPs suggest that regulatory variants constitute a key component of the genetic architecture of grain yield in upland rice. Characterization of these regions revealed an enrichment of *cis*-regulatory elements recognized by transcription factors belonging to the bZIP, MYB, bHLH, WRKY, NAC, ARF, GAMYB, and ERF families—factors involved in coordinating processes related to hormonal signaling, carbon metabolism, vegetative and reproductive development, and responses to environmental stresses (Meraj et al., 2020). Notable among the identified elements are ABRE, MBS, MYC, DRE, W-box, G-box, GARE/P-box, and ERE, which form an integrated regulatory network that simultaneously coordinates growth, development, and grain yield.

**Table 2.** Intergenic SNPs associated with grain yield in upland rice, adjacent candidate genes and the distribution of *cis*-regulatory elements identified within the corresponding genomic regions.

| SNP | Alleles * | Adjacent genes | ABRE | MBS | MYC | DRE | W-box | G-box | GARE/P-box | ERE |
| --- | --- | --- | --- | --- | --- | --- | --- | --- | --- | --- |
| S1_31329438 | C/G | LOC_Os01g54460; LOC_Os01g54470 | X | X | X | X |  | X | X |  |
| S1_22705540 | T/A | LOC_Os01g40230; LOC_Os01g40240 | X | X |  | X | X | X |  |  |
| S2_31420405 | C/G | LOC_Os02g51310; LOC_Os02g51320 | X | X | X |  | X | X | X | X |
| S2_35075916 | G/C | LOC_Os02g57250; LOC_Os02g57260 |  | X | X | X | X | X | X | X |
| S3_4047832 | C/T | LOC_Os03g07920; LOC_Os03g07930 | X | X | X | X | X | X | X | X |
| S3_14490495 | G/A | LOC_Os03g25340; LOC_Os03g25350 | X | X | X | X | X | X | X | X |
| S3_4116916 | C/T | LOC_Os03g08050; LOC_Os03g08070 | X | X | X | X | X | X | X | X |
| S4_28948553 | A/C | LOC_Os04g48560; LOC_Os04g48570 | X | X | X | X |  | X | X |  |
| S5_28442567 | T/G | LOC_Os05g49570; LOC_Os05g49580 | X | X | X | X | X | X | X | X |
| S6_9605790 | A/G | LOC_Os06g16670; LOC_Os06g16680 | X | X | X | X | X | X | X | X |
| S7_18821080 | G/T | LOC_Os07g31690; LOC_Os07g31700 | X | X | X | X | X | X |  | X |
| S8_5699891 | C/A | LOC_Os08g09870; LOC_Os08g09880 | X | X | X |  | X | X |  |  |
| S12_3398630 | C/T | LOC_Os12g06960; LOC_Os12g06970 | X | X | X |  | X | X |  |  |
| Occurrence | - | - | 12 | 13 | 12 | 10 | 11 | 12 | 9 | 8 |

\* Alleles are presented as Reference allele/Alternative allele.

The ABRE, MBS, MYC, and DRE elements highlight the importance of pathways regulated by ABA, MYB, bHLH, and DREB in adaptation to water deficit—a factor particularly relevant for upland cropping systems (Marand et al., 2023). Simultaneously, the high frequency of W-box and G-box elements suggests the involvement of regulatory networks associated with light perception, photosynthesis, carbon assimilation, and energy metabolism—processes that determine the availability of photoassimilates for grain filling (Ma et al., 2023; Huo et al., 2024). Likewise, GARE/P-box and ERE elements indicate the integrated action of auxins, gibberellins, and ethylene in regulating tillering, panicle development, spikelet fertility, and grain filling, reinforcing the concept that grain yield results from the interaction of multiple hormonal and metabolic pathways (Deveshwar et al., 2020; Shaw et al., 2023). These results suggest that the identified regulatory variants can quantitatively modulate gene expression through various mechanisms—including alterations in transcription factor binding sites, linkage disequilibrium with functional regulatory variants, and three-dimensional chromatin interactions—thereby generating cumulative effects on grain yield (Ricci et al., 2019; Marand et al., 2023).

This interpretation is reinforced by the functional convergence between candidate genes identified via GWAS and *cis*-elements located in intergenic regions. Rather than representing independent mechanisms, these components integrate gene regulatory networks that coordinate the perception of environmental stimuli, hormonal signaling, energy metabolism, metabolite transport, and reproductive development. From this perspective, grain yield can be interpreted as an emergent property of these regulatory networks, in which coding and regulatory variants act in concert to modulate physiological processes associated with grain yield (Marand et al., 2023; Huo et al., 2024).

Analysis of the regulatory regions also enabled the establishment of a functional prioritization criterion for the intergenic SNPs identified via GWAS. Although all thirteen SNPs are located in regions enriched with *cis*-regulatory elements, seven variants lie immediately adjacent to—or within a few nucleotides of—these elements, thereby increasing their biological plausibility as potential modulators of transcriptional activity (Table 3). Notable among the identified elements are STRE, as-1, and the CAAT-box, recognized for their roles in environmental stress responses, hormonal signaling integration, and basal transcriptional regulation, respectively (Lescot et al., 2002; Marand et al., 2023). Integrating statistical association, regulatory region characterization, and the presence of biologically plausible candidate genes establishes a consistent hierarchy of evidence for the functional prioritization of these variants, identifying the seven SNPs as the most promising candidates for studies involving gene expression, eQTL analysis, promoter activity, and precision gene editing.

**Table 3.**
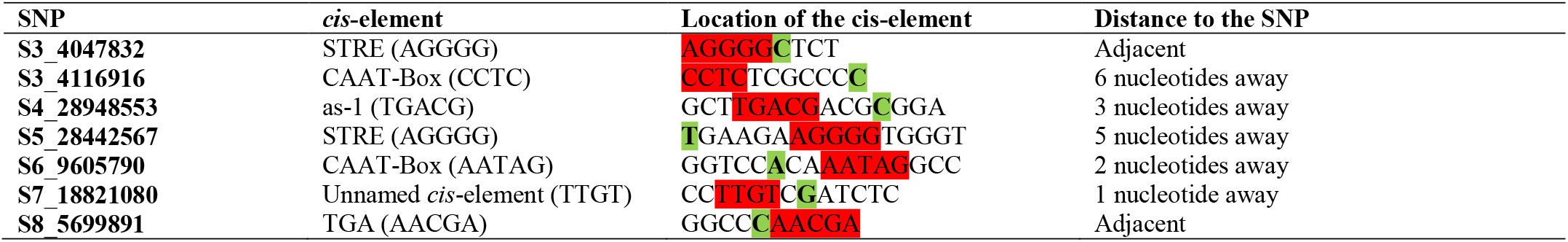
Prioritized intergenic SNPs associated with grain yield in upland rice, their nearest *cis*-regulatory elements, the corresponding regulatory sequences and the physical distance between each SNP and its associated *cis*-element.

Collectively, the results demonstrate that integrating genomic association, functional annotation, and regulatory genomics significantly expands the interpretive potential of GWAS studies, enabling the prioritization of variants with a higher likelihood of biological relevance and applicability to molecular breeding. Beyond identifying new loci associated with grain yield, this study proposes an integrative framework for the discovery, prioritization, and functional validation of regulatory variants; this approach bridges association genomics and functional genomics, providing a conceptual framework applicable to the investigation of other complex quantitative traits in rice and various crop species (Marand et al., 2023; Huo et al., 2024; Springer & Schmitz, 2017).

## CONCLUSIONS

1. This study identified SNPs and candidate genes associated with grain yield in upland rice, expanding the understanding of the genetic architecture of this quantitative trait and demonstrating that coding and regulatory variants act complementarily to determine yield.
2. The integration of GWAS, functional annotation, and regulatory genomics enabled the prioritization of variants with high biological plausibility, revealing that SNPs located in intergenic regions enriched with *cis*-regulatory elements were key candidates for functional validation studies.
3. The results established an integrative framework for the discovery, prioritization, and functional validation of regulatory variants, bridging the gap between association genomics and practical application in molecular breeding, and providing a general framework potentially applicable to the investigation of other complex quantitative traits in rice and various crop species.

## ACKNOWLEDGMENTS

To the National Council for Scientific and Technological Development (CNPq) for financial support (grant no. 407818/2023-5); to the Goiás State Research Foundation (FAPEG) for the scholarship awarded to the fifth author (grant no. 202510267001595); and to the Coordination for the Improvement of Higher Education Personnel (CAPES, Finance Code 001) for the scholarship awarded to the first author.

## REFERENCES

Ashfaq, M.; Rasheed, A.; Zhu, R.; Ali, M.; Javed, M.A.; Anwar, A.; Tabassum, J.; Shaheen, S.; Wu, X. Genome-wide association mapping for yield and yield-related traits in rice (Oryza sativa L.) using SNPs markers. Genes, 14: 1089, 2023. 10.3390/genes14051089

Bradbury, P.J.; Zhang, Z.; Kroon, D.E.; Casstevens, T.M.; Ramdoss, Y.; Buckler, E.S. (2007) TASSEL: software for association mapping of complex traits in diverse samples. Bioinformatics, 19:2633–2635, 2007.10.1093/bioinformatics/btm308

Deveshwar, P.; Prusty, A.; Sharma, S.; Tyagi, A.K. Phytohormone-mediated molecular mechanisms involving multiple genes and QTL govern grain number in rice. Frontiers in Genetics, 11:586462, 2020. 10.3389/fgene.2020.586462

Earl, D.A.; Von Holdt, B.M. STRUCTURE HARVESTER: a website and program for visualizing STRUCTURE output and implementing the Evanno method. Conservation Genetics Resources, 4:359–361, 2012. 10.1007/s12686-011-9548-7

Evanno, G.; Regnaut, S.; Goudet, J. Detecting the number of clusters of individuals using the software STRUCTURE: a simulation study. Molecular Ecology, 14:2611–2620, 2005. 10.1111/j.1365-294X.2005.02553.x

Fernando, Y.; Ovenden, B.; Sreenivasulu, N.; Butardo, V., Climate adaptation strategies for maintaining rice grain quality in temperate regions. Biology, 14:801, 2025. 10.3390/biology14070801

Heng, Y.; Wu, C.; Long, Y.; Luo, S. et al. OsALMT7 maintains panicle size and grain yield in rice by mediating malate transport. The Plant Cell, 30:889–906, 10.1105/tpc.17.00998

Hu, P.; Wen, Y.; Wang, Y.; Wu, H.; Wang, J.; Wu, K.; Chai, B.; Zhu, L.; Zhang, G.; Gao, Z.; et al. Identification and characterization of Short Crown Root 8, a temperature-sensitive mutant associated with crown root development in rice. International Journal of Molecular Sciences, 22:9868, 2021.10.3390/ijms22189868

Huang, P.; Gu, Q.; Hu, Y.; Li, H.; Wu, Z.; Liu, W.; Zhu, Z.; Yuan, P.; Duan, L.; Zhou, Y.; et al. Genetic analysis of a collection of rice germplasm (Oryza sativa L.) through high-density snp array provides useful information for further breeding practices. Genes, 13:830, 2022. 10.3390/genes13050830

Huo, Q.; Song, R.; Ma, Z. Recent advances in exploring transcriptional regulatory landscape of crops. Frontiers in Plant Sciences, 15:1421503, 2024.10.3389/fpls.2024.1421503

Jain, M.; Nijhawan, A.; Arora, R.; Agarwal, P.; Ray, S.; Sharma, P.; Kapoor, S.; Tyagi, A.K.; Khurana, J.P. F-box proteins in rice. Genome-wide analysis, classification, temporal and spatial gene expression during panicle and seed development, and regulation by light and abiotic stress. Plant Physiology, 143:1467–1483, 2007. 10.1104/pp.106.091900

Joly-Lopez, Z., Platts, A.E., Gulko, B. et al. An inferred fitness consequence map of the rice genome. Nature Plants, 6:119–130, 2020. 10.1038/s41477-019-0589-3

Lescot, M.; Déhais, P.; Thijs, G.; Marchal, K.; Moreau, Y.; Van De Peer, Y.; Rouzé, P.; Rombauts, S. PlantCARE, a database of plant cis-acting regulatory elements and a portal to tools for in silico analysis of promoter sequences. Nucleic Acids Research, 30:325–327, 2002. 10.1093/nar/30.1.325

Li, X.; Lappalainen, T.; Bussemaker, H.J. Identifying genetic regulatory variants that affect transcription factor activity. BioRxiv, 2022. 10.1101/2022.10.21.513166

Liu, G.; Wang, H.; Gao, H.; Yu, S.; Liu, C.; Wang, Y.; Sun, Y.; Zhang, D. Alternative splicing of functional genes in plant growth, development, and stress responses. International Journal of Molecular Sciences, 26:5864, 2025. 10.3390/ijms26125864

Ma, B.; Zhang, L.; He, Z. Understanding the regulation of cereal grain filling: the way forward. Journal of Integrative Plant Biology, 65:526–547, 2023. 10.1111/jipb.13456

Manzo, G.; Borkowski, K.; Ovcharenko, I. Comparative analysis of deep learning models for predicting causative regulatory variants. Genes, 16:1223, 2025. 10.3390/genes16101223

Marand, A.P.; Eveland, A.L.; Kaufmann, K.; Springer, N. cis-Regulatory elements in plant development, adaptation, and evolution. Annual Review of Plant Biology, 74:111–137, 2023. 10.1146/annurev-arplant-070122-030236

Meng, L.; Du, M.; Zhu, T.; Li, G.; Ding, Y.; Zhang, Q. (2024) PPR proteins in plants: roles, mechanisms, and prospects for rice research. Frontiers in Plant Sciences, 15:1416742, 2024. 10.3389/fpls.2024.1416742

Meraj, T.A.; Fu, J.; Raza, M.A.; Zhu, C.; Shen, Q.; Xu, D.; Wang, Q. Transcriptional factors regulate plant stress responses through mediating secondary metabolism. Genes, 11:346, 2020. 10.3390/genes11040346

Nagasawa, N.; Hibara, K.I.; Heppard, E.P.; Velden, K.A.V.; Luck, S.; Beatty, M.; Nagato, Y.; Sakai, H. GIANT EMBRYO encodes CYP78A13, required for proper size balance between embryo and endosperm in rice. The Plant Journal, 75:592–605, 2013. 10.1111/tpj.12223

Pantalião, G.F.; Narciso, M.; Guimarães, C.; Castro, A.; Colombari, J. M.; Breseghello, F.; Rodrigues, L.; Vianello, R. P.; Borba, T. O.; Brondani, C. Genome wide association study (GWAS) for grain yield in rice cultivated under water deficit. Genetica, 144:651–664, 2016.10.1007/s10709-016-9932-z

Peng, B.; Liu, Y.; Qiu, J.; Peng, J.; Sun, X.; Tian, X.; Zhang, Z.; Huang, Y.; Pang, R.; Zhou, W.; Zhao, J.; Sun, Y.; Wang, Q. (2024) OsG6PGH1 affects various grain quality traits and participates in the salt stress response of rice. Frontiers Plant Sciences, 15:1436998, 2024. 10.3389/fpls.2024.1436998

Pritchard, J.K., Stephens, M., Rosenberg, N.A.; Donnelly, P. Association mapping in structured populations. The American Journal of Human Genetics, 67:170–181, 2000. 10.1086/302959

Qi, T.; Song, L.; Guo, Y.; Chen, C.; Yang, J. From genetic associations to genes: methods, applications, and challenges. Trends in Genetics, 40:642–667, 2024. 10.1016/j.tig.2024.04.008

Ricci, W.A.; Lu, Z.; Ji, L.; Marand, A.P. et al. Widespread long-range cis-regulatory elements in the maize genome. Nature Plants, 5:1237–1249, 2019. 10.1038/s41477-019-0547-0

Shaw, B.P.; Sekhar, S.; Panda, B.B.; Sahu, G.; Chandra, T.; Parida, A.K. Grain filling in rice vis-a-vis ethylene production in the spikelets. Crop Science, 63:351–361, 2023. 10.1002/csc2.20910

Singh, A.K.; Kumar, D.; Gemmati, D.; Ellur, R.K. et al. Investigating genetic diversity and population structure in rice breeding from association mapping of 116 accessions using 64 polymorphic SSR markers. Crops, 4:180–194, 2024. 10.3390/crops4020014

Song, S.; Chen, Y.; Liu, L.; Wang, Y.; Bao, S.; Zhou, X.; Teo, Z.W.; Mao, C.; Gan, Y.; Yu, H. OsFTIP1-Mediated regulation of florigen transport in rice is negatively regulated by the ubiquitin-like domain kinase OsUbDKγ4. Plant Cell, 29:491–507, 2017. 10.1105/tpc.16.00728

Springer, N.M.; Schmitz, R.J. Exploiting induced and natural epigenetic variation for crop improvement. Nature Review Genetics, 18:563–575, 2017. 10.1038/nrg.2017.45

Storey, J.D. A direct approach to false discovery rates, Journal of the Royal Statistical Society, Series B, 64:479–498, 2002. 10.1111/1467-9868.00346

Wang, S.L.; Zhang, Z.H.; Fan, Y.Y. et al. Control of grain weight and size in rice (Oryza sativa L.) by OsPUB3 Encoding a U-Box E3 Ubiquitin Ligase. Rice, 15:58, 2022. 10.1186/s12284-022-00604-1

Xu, F.; Fang, J.; Ou, S.; Gao, S.; Zhang, F.; Du, L.; Xiao, Y.; Wang, H.; Sun, X.; Chu, J.; Wang, G.; Chu, C. Variations in CYP78A13 coding region influence grain size and yield in rice. Plant Cell Environment, 38:800–11, 2015. 10.1111/pce.12452

Zhu, T.; Xia, C.; Yu, R.; Zhou, X.; Xu, X.; Wang, L.; Zong, Z.; Yang, J.; Liu, Y.; Ming, L.; You, Y.; Chen, D.; Xie, W. Comprehensive mapping and modelling of the rice regulome landscape unveils the regulatory architecture underlying complex traits. Nature Communications, 15:6562, 2024. 10.1038/s420141467-024-50787-y

Zou, X.; Qin, Z.; Zhang, C.; Liu, B.; Liu, J.; Zhang, C.; Lin, C.; Li, H.; Zhao, T. Over-expression of an S-domain receptor-like kinase extracellular domain improves panicle architecture and grain yield in rice. Journal of Experimental Botany, 66:7197–7209, 2015. 10.1093/jxb/erv417

